# Accumulation of auranofin in white adipose tissues lowers leptin levels and exerts anti-diabetic effects

**DOI:** 10.1101/2021.09.11.459899

**Authors:** Aaron R. Cox, Peter M. Masschelin, Pradip K Saha, Jessica B. Felix, Robert Sharp, Zeqin Lian, Yan Xia, Natasha Chernis, David A. Bader, Kang Ho Kim, Xin Li, Jun Yoshino, Xin Li, Zheng Sun, Huaizhu Wu, Cristian Coarfa, David D. Moore, Samuel Klein, Kai Sun, Sean M. Hartig

## Abstract

Low-grade, sustained inflammation in white adipose tissue (WAT) characterizes obesity and frequently coincides with insulin resistance and type 2 diabetes (T2D). However, pharmacological targeting of WAT inflammation lacks durable therapeutic effects. Through a computational screen, we identified the FDA-approved rheumatoid arthritis drug auranofin is a putative small molecule for obesity treatment. We discovered that allometrically scaled safe auranofin doses homed to WAT and improved insulin sensitivity in obese wild-type mice. Auranofin treatment also normalized other obesity-associated abnormalities, including hepatic steatosis and hyperinsulinemia. Surprisingly, the anti-diabetic effects of auranofin required leptin lowering and beta-adrenergic receptors in WAT. These metabolic benefits of leptin reduction were superior to any immune impacts of auranofin in WAT. Our studies reveal important metabolic properties of anti-inflammatory treatments and contribute to the notion that leptin reduction in the periphery can be accomplished to treat obesity and T2D.

## INTRODUCTION

The obesity epidemic contributes to the increased health burden of chronic inflammatory conditions, including insulin resistance, type 2 diabetes mellitus (T2DM), fatty liver, and cardiovascular disease (Afshin et al., 2017). Obesity reflects facultative white adipose tissue (WAT) expansion that occurs during prolonged dietary stress. Although some clinical relationships explain how excess body weight causes insulin resistance in most individuals (Smith et al., 2019), WAT inflammation remains among the most debated incident condition associated with obesity and its co-morbidities.

Obesity promotes chronic low-grade inflammation in WAT that governs local and systemic metabolic responses to overnutrition (Saltiel and Olefsky, 2017). Along these lines, many studies demonstrate WAT inflammation causes local and systemic insulin resistance in rodents (Kanda et al., 2006; Lee et al., 2011; Lumeng et al., 2007; Patsouris et al., 2008; Weisberg et al., 2003) and widely used anti-diabetic drugs exert anti-inflammatory effects as part of their therapeutic activity (Isoda et al., 2006; Soccio et al., 2014). Although WAT inflammation is one conserved pathway that links obesity to insulin resistance and T2DM, directly targeting the immune component of obesity has proved elusive, and broad anti-inflammatory strategies lack clinical efficacy in the treatment of obesity (Goldfine et al., 2013; Goldfine et al., 2010; Ofei et al., 1996). Nonetheless, there is an uncontroversial relationship between low-grade inflammation in WAT and the onset of insulin resistance in humans and rodents. This knowledge gap precludes the development of immune therapies to treat the pressing medical problem of obesity and its metabolic complications, including insulin resistance, fatty liver disease, and T2DM.

We previously demonstrated enforced expression of the microRNA *miR-30a* in subcutaneous fat tissues caused anti-diabetic effects in mouse models (Koh et al., 2016; Koh et al., 2018; Saha et al., 2020). We further showed *miR-30a* opposed the actions of macrophage and T cell cytokines in WAT, suggesting an important role for *miR-30a* in defending adipocytes against pro-inflammatory signals. Notably, we also reported expression of *miR-30a* in WAT corresponds with insulin sensitivity in obese mice and humans (Koh et al., 2018; Saha et al., 2020). Our findings supported the concept that sustained *miR-30a* expression or surrogate approaches may uncouple obesity from the metabolic consequences of overnutrition.

Effective prevention and therapeutic approaches are needed to lower the prevalence of obesity and T2DM and, ultimately, improve health outcomes. The present study grew from computational methods motivated by the idea that we can repurpose FDA-approved therapies for obesity that generate *miR-30a-*like effects. To this end, we used pre-clinical models to explore novel interventions that might slow the progression of obesity to T2DM among at-risk individuals.

## RESULTS AND DISCUSSION

We hypothesized small molecules that drive immune and metabolic profiles similar to ectopic *miR-30a* (Adv-*miR-30a*) expression in inguinal WAT (iWAT) generate anti-diabetic effects. To explore this hypothesis, we used the Broad Connectivity Map (https://clue.io/cmap) to screen more than 5000 small molecules for mRNA profiles that resemble ectopic *miR-30a* expression in iWAT **(Figure 1A)**. Through this process, we identified auranofin (2,3,4,6-tetra-O-acetyl-1-thio-β-D-glucopyranosato-S-triethylphosphine-gold) as a compound with an identical RNA-Seq profile as Adv-*miR-30a*. In addition to known effects on pro-inflammatory responses (Nakaya et al., 2011), auranofin (Ridaura) was intriguing to us because it is an FDA-approved, orally available gold salt for the treatment of rheumatoid arthritis, an inflammatory condition often coupled with hyperglycemia, insulin resistance, and the comorbidities of T2DM (Giles et al., 2018). In human patients who received gold salt injections, traces of gold can be found in numerous tissues (Gottlieb et al., 1972; Vernon-Roberts et al., 1976). Therefore, we administered auranofin to high-fat diet (HFD)-induced obese mice to define the tissue distribution of auranofin. Surprisingly, mass spectrometry methods indicated that auranofin selectively accumulates in epididymal (eWAT) and inguinal fat compartments **(Figure 1B)**, which suggested efficient and unique drug targeting to the inflammatory environment of WAT in obesity.

**Figure 1.**
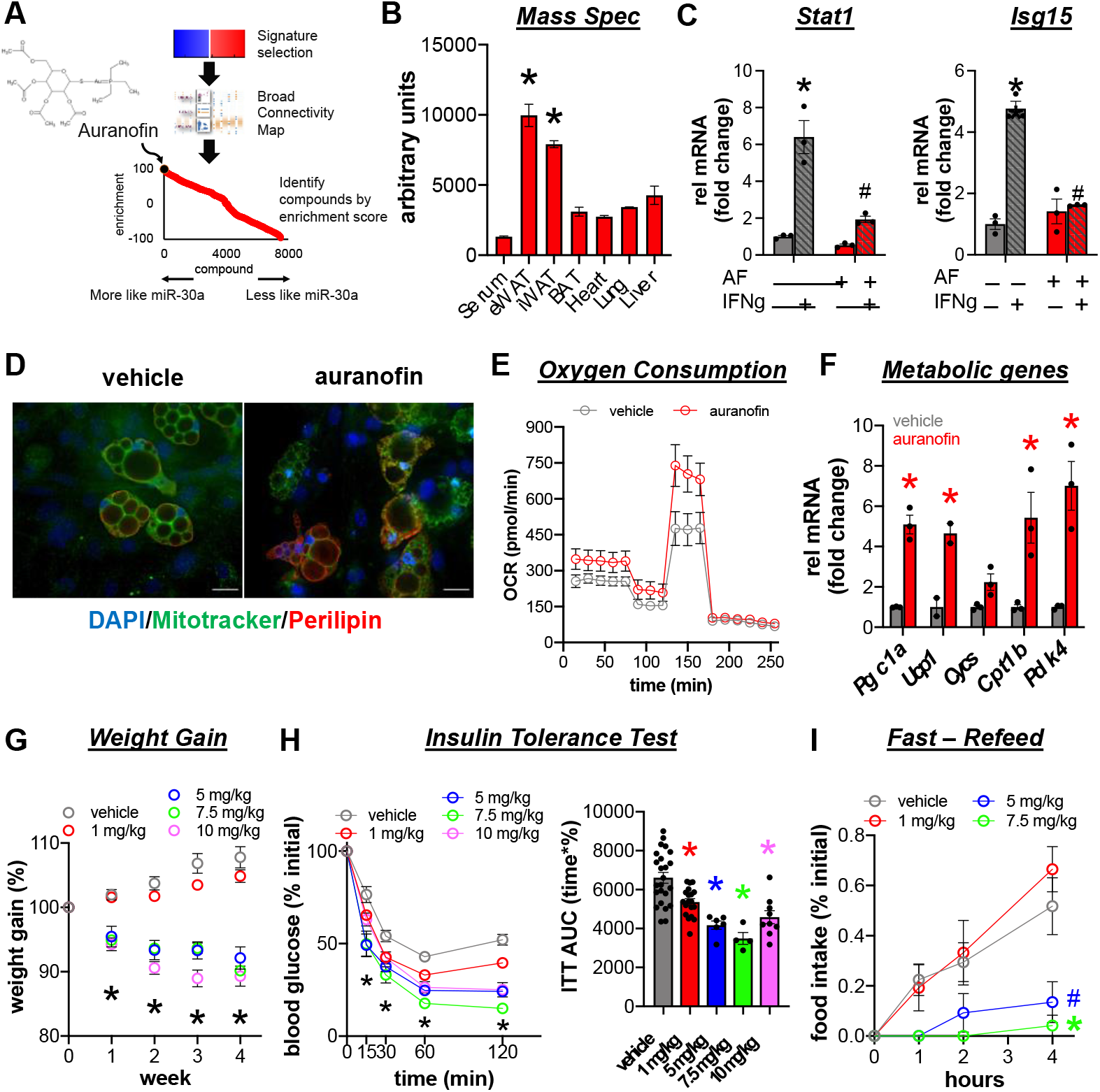
The anti-rheumatic drug auranofin and *miR-30a* exhibit similar activities. **(A)** The Broad connectivity map identified compounds with expression signatures similar to enforced expression of *miR-30a* in the inguinal WAT of obese mice. This in-silico approach identified the anti-inflammatory rheumatoid arthritis drug auranofin regulates gene expression in a similar way as *miR-30a*. **(B)** Diet-induced obese mice were injected with a single dose of auranofin (10 mg/kg ip) and sacrificed 24 hours later for tissue harvest and mass spectrometry analysis of tissue distribution. *p<0.05 vs all other tissues. **(C)** WAT stromal vascular cells (SVF) were differentiated into adipocytes and then exposed to auranofin (1000 nM; red) or vehicle (DMSO; gray) +/-IFNg (100 ng/ml; hatched bars). Relative mRNA expression of inflammatory genes. *p<0.05 vs DMSO vehicle; #p<0.05 vs IFNg alone. **(D)** Immunofluorescence (DAPI – blue, perilipin – red, MitoTracker – green) images of differentiated adipocytes from WAT SVF in the presence of auranofin (100 nM) or DMSO for 24 h. Scale bar, 20 µm. **(E)** Respiration (as oxygen consumption rate, OCR) in differentiated mouse adipocytes measured during the Seahorse XF Mitochondrial Stress kit. **(F)** Relative mRNA expression of metabolic genes in differentiated adipocytes exposed to auranofin (1000 nM; red) or vehicle (DMSO; gray). *p<0.05 vs vehicle. Wild-type mice fed a high fat diet (HFD) for 12 weeks were i.p. injected with various doses of auranofin (1, 5, 7.5, 10 mg/kg) or vehicle for 4 weeks. **(G)** Weight gain (% initial). vehicle n=23, 1 mg/kg n=29, 5 mg/kg n=6, 7.5 mg/kg n=4, 10 mg/kg n=10. *p<0.05 week 1 vehicle vs 7.5, 10 mg/kg and weeks 2-4 vehicle vs 5, 7.5, 10 mg/kg. **(H)** Insulin tolerance tests (ITT) with corresponding area under the curve measurements. vehicle n=24, 1 mg/kg n=19, 5 mg/kg n=6, 7.5 mg/kg n=4, 10 mg/kg n=9. *p<0.05 vehicle vs all doses. **(I)** Food intake (% initial) was measured in mice fasted overnight for 16 h and then injected with auranofin followed by re-feeding (n=4-5/group). *p<0.05 vehicle vs 7.5 mg/kg; #p<0.08 vehicle vs 5 mg/kg. Data are represented as mean ± SEM.

Auranofin exerts diverse effects in many different cell types in vitro (Balfourier et al., 2020). To examine the cell-autonomous impacts of auranofin, we used differentiated adipocytes generated from the iWAT stromal vascular fraction (SVF) of wild-type mice. First, we modeled obesity-induced inflammation in WAT by treating differentiated SVF-derived adipocytes with the inflammatory cytokine interferon gamma (IFNγ) for 24 h (Wentworth et al., 2017). Chronic IFNγ treatment increased the mRNA levels of key pro-inflammatory targets, *Stat1* and *Isg15* **(Figure 1C)**. These effects were prevented entirely by co-treatment with auranofin. Auranofin-treated cells also exhibited more densely packed mitochondria **(Figure 1D)**, contributing to greater respiratory capacity **(Figure 1E)**. Likewise, auranofin heightened the expression of integral mitochondrial and metabolic genes *Ucp1, Pgc1a, Cycs, Cpt1b*, and *Pdk4* that reflect brown and beige adipocyte functions **(Figure 1F)**. These previously unknown drug impacts demonstrate auranofin can block inflammatory signals and empower metabolic functions known to improve insulin sensitivity.

We performed a series of dose escalation studies in obese mice to identify a safe, tolerable treatment regimen. Wild-type mice fed HFD for 12 weeks were i.p. injected with various doses of auranofin three days per week (Monday/Wednesday/Friday) for one month and then administered an insulin tolerance test. We learned the published treatment regimens (Fiskus et al., 2014; Yan et al., 2019), and two lower doses cause unreported distress, malaise, and dramatic weight loss **(Figure 1G)**. Insulin tolerance tests showed significantly greater responsiveness to insulin in all auranofin-treated groups compared to vehicle controls **(Figure 1H)**. To address the possibility that weight loss derived from hypophagia, we performed a fast-refeeding experiment. After an overnight fast, mice were injected with auranofin or vehicle thirty minutes before refeeding **(Figure 1I)**. Control mice responded immediately to food reintroduction and continued to eat during the 4 h follow-up period. In contrast, mice receiving higher doses of auranofin showed food aversion and exhibited noticeable malaise, suggesting high doses of auranofin cause toxicity. Mice tolerated the 1 mg/kg dose well, and we observed no effects on food intake or gross behaviors. Importantly, allometric scaling indicated the 1 mg/kg dose three days/week is well below human clinical doses (maximum 9 mg/day). These data demonstrate safe doses of auranofin administered at levels ten-fold lower than previous studies (Fiskus et al., 2014; Yan et al., 2019), and about half the bioequivalent in humans improves insulin sensitivity and furnishes previously unrecognized therapeutic benefits in obesity.

With the ultimate goal of repurposing drugs to treat T2DM, we built upon these studies at the 1 mg/kg dose to establish whether auranofin improves the inflammatory and metabolic profile of obesity. Consistent with nominal body weight effects **(Figure 1G)**, magnetic resonance imaging (MRI) measurements of fat and lean mass established body composition of auranofin and vehicle groups were equivalent **(Figure 2A)**. Using indirect calorimetry, we found that auranofin did not significantly change food intake **(Figure 2B)** or respiratory exchange ratio **(Figure 2C)**. However, auranofin treatment in obese mice led to elevated physical activity in the dark cycle **(Figure 2D)**.

**Figure 2.**
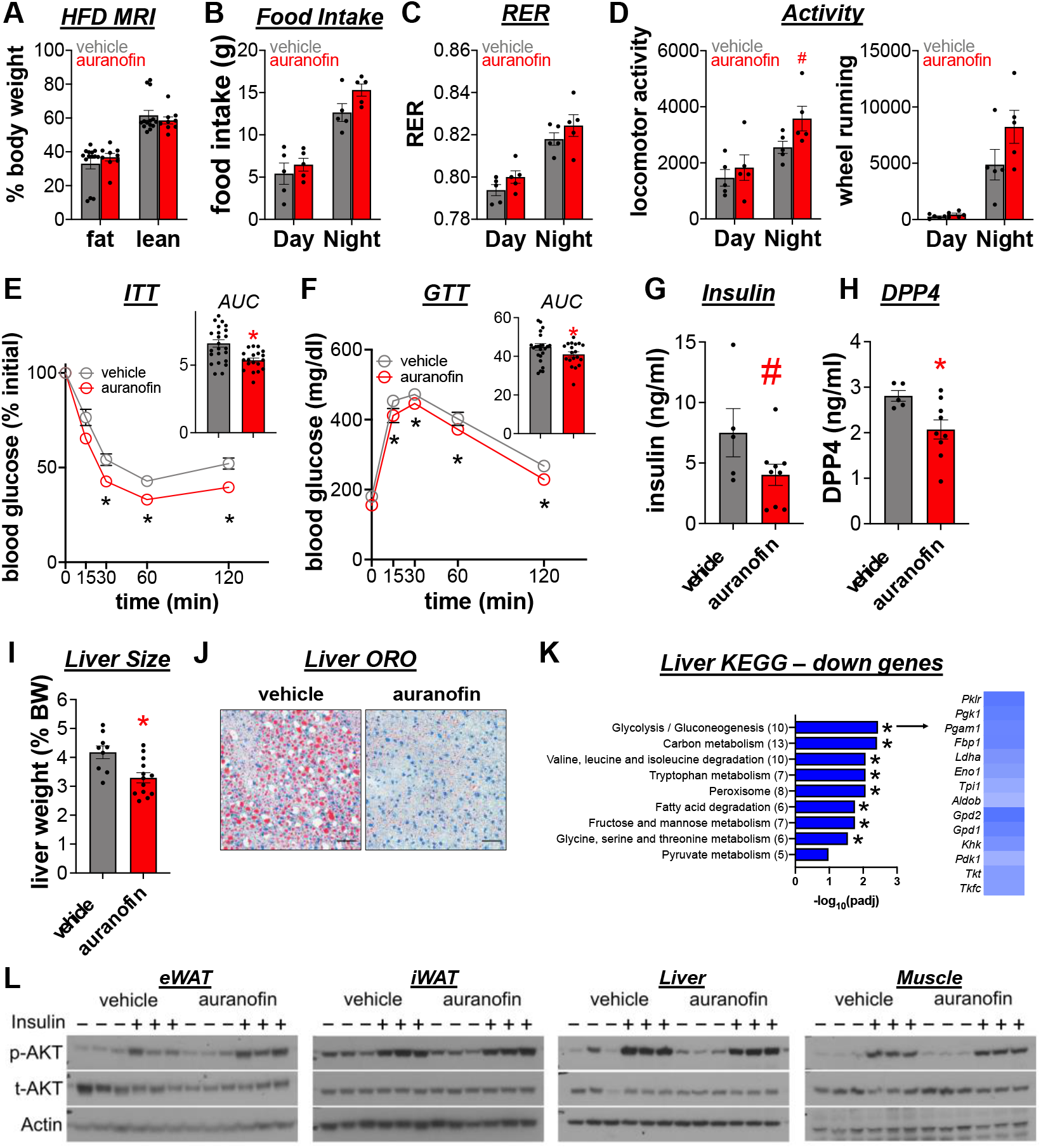
Auranofin improves insulin sensitivity in obese mice. Mice fed a high fat diet (HFD) for 12 weeks were i.p. injected with auranofin (1 mg/kg; red) or vehicle (gray) for 4 weeks. **(A)** Body composition (% body mass; n=9-14/group). Mice were individually housed and monitored in CLAMS home cages for 7 days (n=5). Averaged data during dark and light periods for **(B)** food intake **(C)** RER and **(D)** activity level measured by locomotor and wheel running. *p<0.05 vs vehicle; #p<0.07 vs vehicle. **(E)** Insulin (ITT; n=19-24/group) and **(F)** glucose (GTT; n=19-24/group) tolerance tests with corresponding area under the curve measurements, followed by **(G)** fasting serum insulin (ng/ml; n=5-9/group) and **(H)** fed serum DPP4 levels. *p<0.05 vs vehicle, #p<0.09 vs vehicle. **(I)** Liver size (% BW; n=9-13/group) and **(J)** Oil Red O (ORO) staining of liver sections from HFD mice treated with vehicle or auranofin (1 mg/kg). Scale bar, 100 µm. **(K)** KEGG pathway analysis of down-regulated genes by RNA-seq in the liver of HFD obese mice treated with auranofin shown as –log_10_(adjusted p-value). Heatmap (right) shows RPKM values for down-regulated genes enriched in glycolysis/gluconeogenesis, as well as fatty acid metabolism (*Khk, Pdk1, Tkt, Tkfc*). **(L)** Immunoblots of tissue lysates from mice injected with insulin five minutes prior to sacrifice and probed for p-Akt(Ser473) and total Akt with actin as a loading control. Data are represented as mean ± SEM.

We expanded the earlier studies to comprehensively understand the impacts of auranofin on glucose metabolism. Auranofin treatment significantly improved insulin **(Figure 2E)** and glucose tolerance **(Figure 2F)** in obese mice and corrected fasting hyperinsulinemia **(Figure 2G)**. Of note, auranofin reduced serum dipeptidyl-peptidase-4 **(Figure 2H)**, a liver-derived inflammatory protein that circulates at higher levels in diabetic mice and humans (Mulvihill and Drucker, 2014). Consistent with known associations between higher levels of circulating DPP-4 protein and progression of fatty liver disease (Baumeier et al., 2017; Williams et al., 2015), we observed that auranofin reduced liver weight **(Figure 2I)** and hepatic lipid accumulation **(Figure 2J)**. RNA-seq coupled with pathway analysis established auranofin reduced the activity of several metabolic pathways known to cluster with fatty liver disease, including glycolysis and gluconeogenesis **(Figure 2K)**. Additionally, auranofin suppressed the expression of several genes involved in lipogenesis (*Khk, Pdk1, Tkt, Tkfc*). We next asked whether insulin sensitivity changes in auranofin-treated mice reflected improved insulin signaling **(Figure 2L)**. Despite improved fatty liver profiles, auranofin only increased insulin-dependent phosphorylation of Akt at S473 in the epididymal WAT (eWAT). Collectively, these data demonstrate auranofin increases insulin sensitivity in tissue-specific ways and improves the metabolic phenotype of obese mice.

Next, we studied the effect of auranofin on glucose homeostasis in lean mice. Eighteen-week-old wild-type mice on a normal chow diet were given auranofin (1 mg/kg) three days per week (Monday/Wednesday/Friday) for one month, followed by comprehensive metabolic phenotyping. No malaise or body composition effects were observed in lean mice receiving auranofin compared to vehicle controls **(Figure S1A-C)**. GTT **(Figure S1D)** and ITT **(Figure S1E)** performed on lean mice treated with auranofin or vehicle showed no differences in glucose tolerance and insulin sensitivity. Fasting insulin levels were also equivalent between auranofin and control mice **(Figure S1F)**. These data argue auranofin actions require obesity.

During sustained HFD feeding, obesity initiates low-grade metabolic inflammation that ultimately coincides with insulin resistance and T2DM. In this setting, eWAT harbors chronic inflammation and tissue dysfunction (Lumeng et al., 2007; Weisberg et al., 2003). Drug accumulation **(Figure 1B)** and enhanced insulin action in eWAT **(Figure 2L)** suggested auranofin may influence changes in the eWAT compartment that govern local tissue inflammation. Histology **(Figure 3A)** and quantitative image-based analysis revealed adipocytes from auranofin-treated mice were significantly smaller **(Figure 3B)**, favoring reduced adipocyte hypertrophy (Jeffery et al., 2016). We used RNA-Seq to identify the biologically cohesive gene programs of auranofin in the eWAT of obese mice. These efforts uncovered clear signatures that explain the impacts of auranofin in the eWAT. Consistent with smaller adipocytes **(Figure 3C)**, auranofin increased the expression of adipogenesis markers (*Adipoq, Glut4, Acaca, Scd1*) and concurrently depleted levels of preadipocyte genes (*Prrx1, Fsp1*). Moreover, Gene Set Enrichment Analysis (GSEA) indicated auranofin elevated levels of genes found in central metabolic pathways, including oxidative phosphorylation, adipogenesis, and fatty acid metabolism. Not surprisingly, GSEA also demonstrated wide anti-inflammatory effects reflected by changes in marker genes of macrophages, innate immunity, and the inflammasome **(Figure 3E)**.

**Figure 3.**
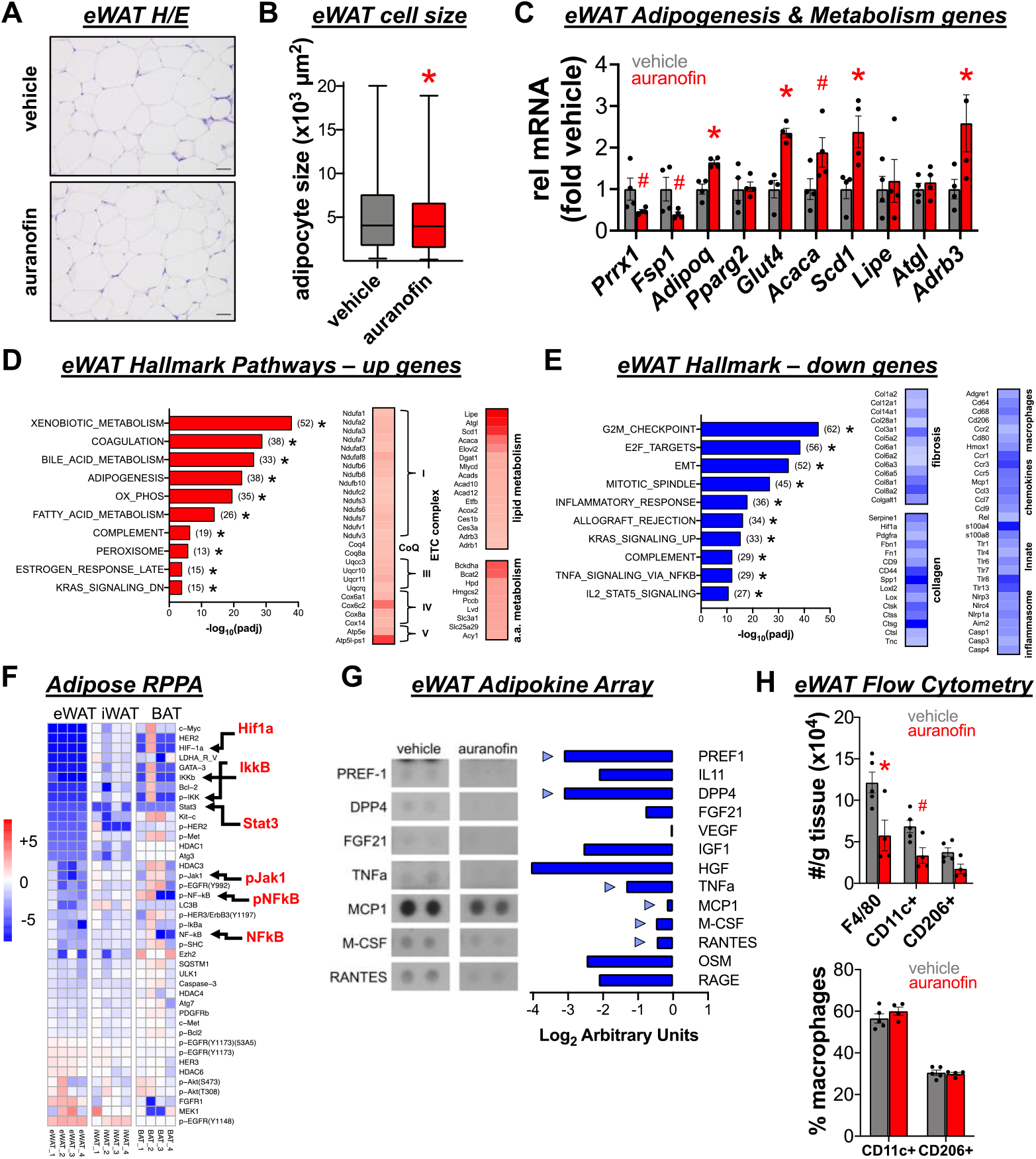
Auranofin suppresses eWAT inflammation in obese mice. Mice fed a high fat diet (HFD) for 12 weeks were i.p. injected with auranofin (1 mg/kg; red) or vehicle (gray) for 4 weeks. **(A)** eWAT H&E and **(B)** mean adipocyte size (µm^2^) tabulated across four 20x fields of view (n=3/group). *p<0.05. **(C)** Relative mRNA expression of adipogenesis and metabolism genes from eWAT of obese mice treated with auranofin (red) or vehicle (gray). *p<0.05 vs vehicle; #p<0.10 vs vehicle. KEGG pathway analysis of **(D)** up-regulated and **(E)** down-regulated genes by RNA-seq in the eWAT of HFD obese mice treated with auranofin shown as – log_10_(adjusted p-value). Heatmaps show RPKM values for **(D)** up-regulated genes enriched in mitochondrial complexes, lipid and amino acid metabolism and **(E)** down-regulated genes enriched in collagen, fibrosis, and inflammation (macrophages, chemokines, innate signaling, and inflammasome). **(F)** Reverse phase protein array on eWAT, iWAT, and BAT of HFD obese mice shown as a heatmap of log_2_(fold change auranofin over vehicle). Arrows highlight pro-inflammatory proteins and phospho-proteins suppressed by auranofin in eWAT. **(G)** eWAT protein lysates (pooled from n=4/group) were incubated with adipokine spotted arrays (left) and quantified for relative abundance as log_2_(fold change auranofin over vehicle) (right). Arrowheads highlight pro-inflammatory cytokines and chemokines. **(H)** Flow cytometry analysis of macrophage populations in eWAT marked by F4/80, CD11c+, and CD206+ expressed as number/g tissue (×10^4^) (top) or % F4/80+ macrophages (bottom) for obese mice treated with auranofin (red) and vehicle (gray). *p<0.05 vs vehicle; #p<0.10 vs vehicle. Data are represented as mean ± SEM.

To further characterize cell signaling events underpinning the metabolic and inflammatory phenotypes, we applied WAT and brown adipose tissue (BAT) samples to a reverse phase protein array (RPPA) with broad pathway coverage **(Figure 3F)**. Auranofin imparted the most robust impacts on eWAT probes, including depletion of well-established protein markers of inflammation in the NF-kB (IKKb, p-IKK, p-IkBa, NF-kB, p-NF-kB) and JAK-STAT (p-JAK1, STAT3) signaling pathways. Blocking these pathways coincided with decreased levels of pro-inflammatory cytokines (PREF1, DPP4, TNFa) and chemokines (MCP1, M-CSF, RANTES) in the eWAT **(Figure 3G)**. These data suggested decreased recruitment and function of macrophages, so we performed flow cytometry on the eWAT SVF. Total macrophages labeled by F4/80 (ADGRE1) were significantly reduced with auranofin treatment, with a strong trend towards fewer total CD11c+ pro-inflammatory M1-like macrophages **(Figure 3H)**. The percent macrophages in each subtype did not change, suggesting across-the-board depletion rather than unique targeting of CD11c+ or CD206+ immune cells. Together, our findings suggest auranofin remodels the eWAT compartment through enhanced adipogenesis and adipocyte metabolism coupled with suppression of tissue macrophage pro-inflammatory competence.

Our previous findings (Koh et al., 2018) demonstrated *miR-30a* improved the inflammatory profile of obesity and formed the foundation to nominate auranofin as a putative anti-diabetic therapy **(Figure 1A)**. Because auranofin performs similar metabolic and anti-inflammatory effects as the ectopic *miR-30a* expression in WAT, we investigated whether *miR-30a* knockout accounted for the health benefits of auranofin in obese mice. To explore whether *miR-30a* expression explained the phenotype of auranofin treatment, we used CRISPR/Cas9 gene editing to knockout the entire *miR-30a* gene **(Figure 4A)**. *miR-30a* knockout was confirmed using qPCR analysis and DNA sequencing. The *miR-30a*^*-/-*^ mice were viable and showed no overt phenotypes during postnatal growth. *miR-30a*^*-/-*^ mice on a HFD exhibited similar body weight **(Figure 4B)**, ad-libitum blood glucose **(Figure 4C)**, and serum insulin levels **(Figure 4C)** compared to wild-type mice.

**Figure 4.**
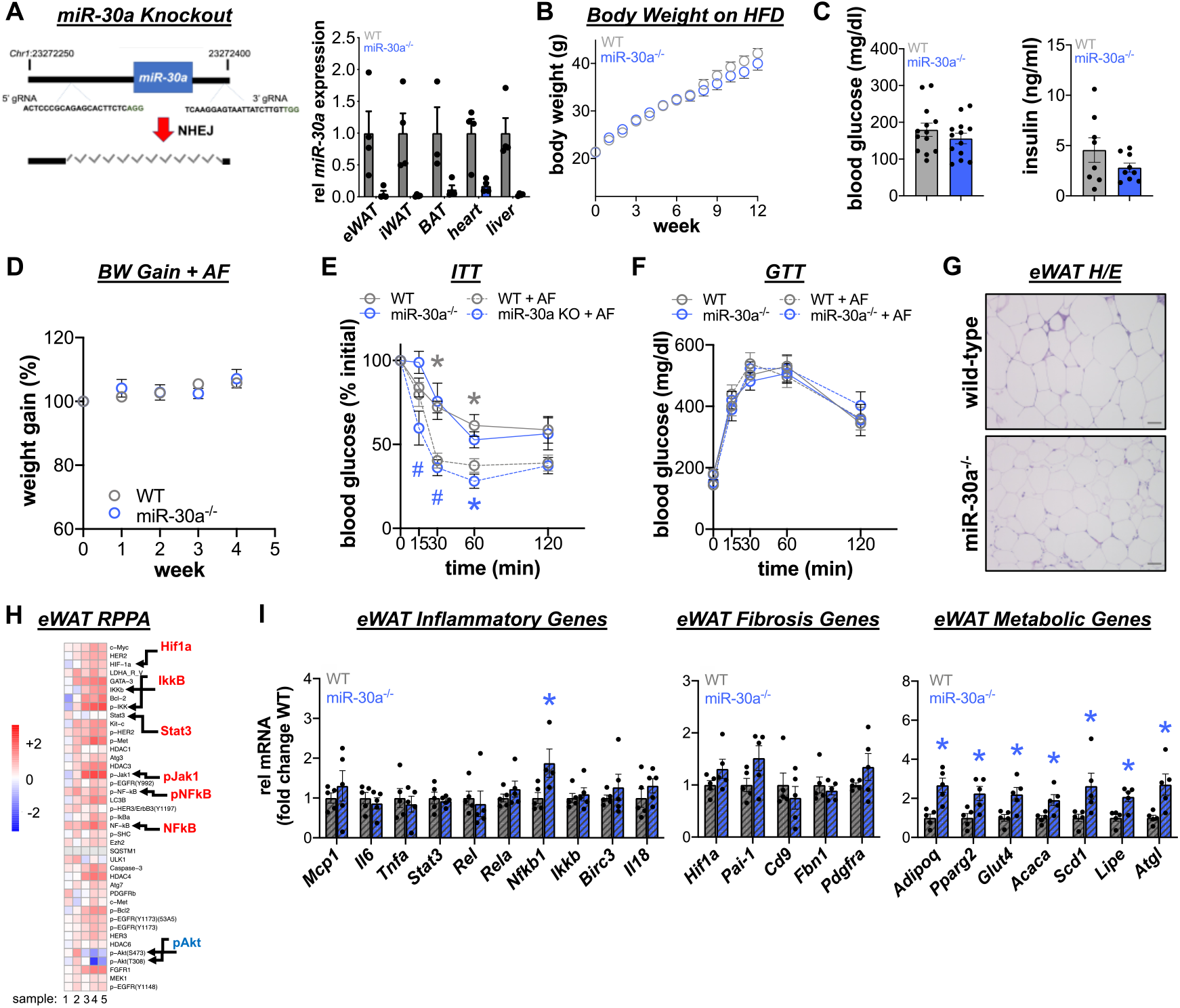
*miR-30a* provides a fractional contribution to the effects of auranofin. **(A)** Guide RNAs (gRNA) flanked a 1 kb region on mouse chromosome 1 that contains *miR-30a*. Non-homologous end-joining (NHEJ) repaired CRISPR-mediated DNA breaks. qPCR analysis of tissues collected from six-month old *miR-30a*^*-/-*^ mice and littermate controls (n=4/group). **(B)** Body weight of wild-type and *miR-30a*^*-/-*^ mice fed a high fat diet (HFD) for 12 weeks (n=21-22 mice/group). **(C)** Fasting glucose (n=12/group) and insulin levels (n=8-9/group). Obese wild-type and *miR-30a*^*-/-*^ mice were i.p. injected with auranofin (1 mg/kg) for 4 weeks. **(D)** Body weight gain of HFD wild-type (gray) and *miR-30a*^*-/-*^ (blue) mice during auranofin treatment. **(E)** Glucose (GTT) and **(F)** insulin (ITT) tolerance tests. Solid lines/bars = before auranofin treatment, dotted lines/hatched bars = after auranofin treatment, wild-type (gray) and *miR-30a*^*-/-*^ (blue). n=5/group; *p<0.05 vs vehicle. **(G)** H&E stained eWAT in HFD wild-type and *miR-30a*^*-/-*^ mice treated with auranofin. **(H)** Reverse phase protein array (RPPA) on eWAT of untreated HFD obese mice shown as a heatmap of log_2_(fold change versus wild-type). **(I)** Relative mRNA expression of inflammatory and fibrosis genes from eWAT of HFD wild-type (gray hatched) and *miR-30a*^*-/-*^ (blue hatched) mice treated with auranofin. *p<0.05 vs wild-type. Data are represented as mean ± SEM.

We performed longitudinal profiling of body weight and insulin sensitivity in wild-type and *miR-30a*^*-/-*^ mice on HFD before and after auranofin treatment. After 12 weeks on HFD, *miR-30a*^*-/-*^ and wild-type mice were treated with auranofin (1 mg/kg) for four weeks. Both genotypes showed nominal weight gain differences **(Figure 4D)**, glucose intolerance **(Figure 4E)**, and insulin resistance **(Figure 4F)** before auranofin treatment. Although glucose tolerance **(Figure 4E)** did not change amongst treatments and genotypes, auranofin significantly improved insulin sensitivity in both *miR-30a*^*-/-*^ and wild-type mice **(Figure 4F)**. The strong influence of *miR-30a* on WAT inflammation prompted us to assess eWAT inflammation and fibrosis following auranofin treatment in *miR-30a*^*-/-*^ mice. Histological analysis of eWAT tissue sections indicated *miR-30a*^*-/-*^ mice contained abundant mononuclear immune cells resembling wild-type mice **(Figure 4G)**, suggesting auranofin exerted similar outcomes in both genotypes. Of note, RPPA revealed *miR-30a* ablation increased expression of pro-inflammatory NF-kB and JAK-STAT probes in eWAT compared to wild-type mice on HFD **(Figure 4H)**, indicating *miR-30a* and auranofin target the same inflammatory pathways. Indeed, we detected increased *Nfkb1* expression in eWAT of *miR-30a*^*-/-*^ mice treated with auranofin **(Figure 4I)**. Other gene expression profiling showed unremarkable changes in inflammatory and fibrosis markers between wild-type and *miR-30a*^*-/-*^ mice that received auranofin **(Figure 4I)**. Overall, these studies indicate that auranofin and *miR-30a* share similar impacts on some inflammatory effects of HFD, but metabolic phenotypes of auranofin do not require *miR-30a* expression.

In efforts to figure out whether auranofin influenced obesity sequalae in other mouse models, we pursued experiments in leptin-deficient backgrounds that do not need high fat diet interventions for insulin resistance. First, we tested inducible leptin receptor knockout mice (*Ubc-Cre;LepR*^*lp/lp*^; *LepR KO*), an obesity model without the complication of uncontrollable diabetes observed in conventional *db/db* mice (Cox et al., 2016). *LepR KO* mice gained ~17-19 g over 4 weeks after tamoxifen administration **(Figure 5A)**, at which point mice were randomized to receive vehicle or auranofin (1 mg/kg). Auranofin and vehicle treated *LepR KO* mice gained weight at similar rates over the 4-week treatment period **(Figure 5A)** with comparable fat and lean mass levels **(Figure 5B)**. In contrast to observations in obese wild-type mice **(Figure 2)**, *LepR KO* mice resisted the metabolic effects of auranofin. Insulin **(Figure 5C)** and glucose tolerance **(Figure 5D)** tests, as well as fasting serum insulin **(Figure 5E)**, were equivalent in vehicle and auranofin treated *LepR KO* mice. We next assessed the impact of auranofin on inflammation in *LepR KO* mice. Histological analysis of H/E stained eWAT sections **(Figure 5F)**, along with targeted gene expression profiling **(Figure 5G)**, further suggested *LepR KO* do not respond to auranofin. RPPA profiling of eWAT also demonstrated protein probes sensitive to auranofin in wild-type mice unaltered in the *LepR* knockout background. We additionally tested auranofin in leptin-deficient (*ob/ob*) mice and detected similar resistance to the metabolic effects of auranofin **(Figure S2)**. Our enrollment of additional mouse models argued intact leptin signaling furnishes the insulin-sensitizing effects of auranofin.

**Figure 5.**
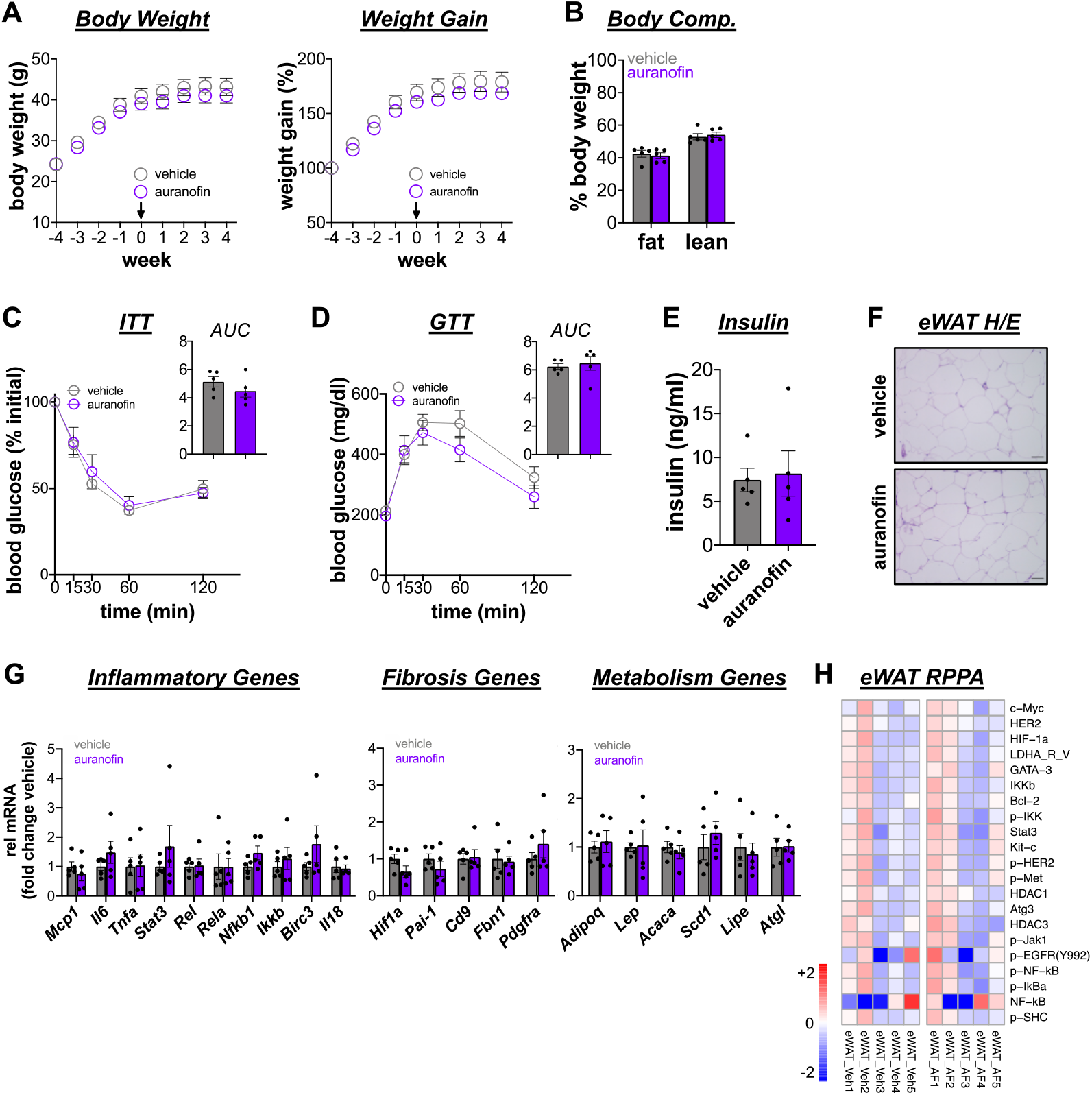
Leptin receptor deficient mice exhibit resistance to the insulin sensitizing effects of auranofin. *Ubc-Cre;LepR*^*lp/lp*^ (*LepR KO*) mice were treated with tamoxifen to induce LepR deletion and obesity. After 4 weeks, *LepR KO* mice were i.p. injected with auranofin (1 mg/kg; purple) or vehicle (gray) for 4 weeks (n=5/group). **(A)** Body weight and weight gain of *LepR KO* during treatment with auranofin or vehicle (n=5/group). **(B)** Final body composition (% body mass). **(C)** Insulin (ITT) and **(D)** glucose (GTT) tolerance tests, with corresponding area under the curve (AUC) measurements (×10^4^ or ×10^5^, respectively). **(E)** Fasting serum insulin (ng/ml). **(F)** H&E stained eWAT. **(G)** Relative mRNA expression of inflammatory, fibrosis, and metabolism genes from eWAT. **(H)** Reverse phase protein array on eWAT shown as a heatmap of log_2_(fold change auranofin over vehicle). Data are represented as mean ± SEM.

Recent studies indicate leptin reduction improves the metabolic phenotype of obesity (Zhao et al., 2020; Zhao et al., 2019). Leptin also induces lipolysis through activation of beta-adrenergic signaling in WAT, a feature largely diminished in chronic obesity and T2DM (Pirzgalska et al., 2017; Zeng et al., 2015; Zhao et al., 2018). We revisited our RNA-seq data and found auranofin significantly increased lipolytic gene expression in the eWAT, including heightened levels of *Adrb3, Adrb1, Atgl, Lipe/Hsl* **(Figure 3D)**. To follow up on these observations, we performed a series of experiments to test the impact of auranofin on the leptin-Adrb3 axis. First, we confirmed auranofin caused significant decreases in serum leptin in HFD-fed wild-type mice compared to controls **(Figure 6A)**, a phenotype consistent with the beneficial effects of partial leptin reduction in obesity (Zhao et al., 2019). In contrast, we observed profound hyperleptinemia in *LepR KO* mice independent of auranofin treatment. Moreover, auranofin significantly induced higher expression of *Adrb3* in wild-type, but not *LepR KO* eWAT **(Figure 6B)**. These data suggest the lipolytic response of eWAT may be impaired in the absence of intact leptin signaling. Importantly, WAT from metabolically unhealthy obese individuals (Cifarelli et al., 2020) exhibit lower *Adrb3* and *Lipe/Hsl* expression than healthy lean controls **(Figure 6C)**, strengthening the notion that restoration of the leptin-Adrb3 axis forms a contemporary method to treat obesity.

**Figure 6.**
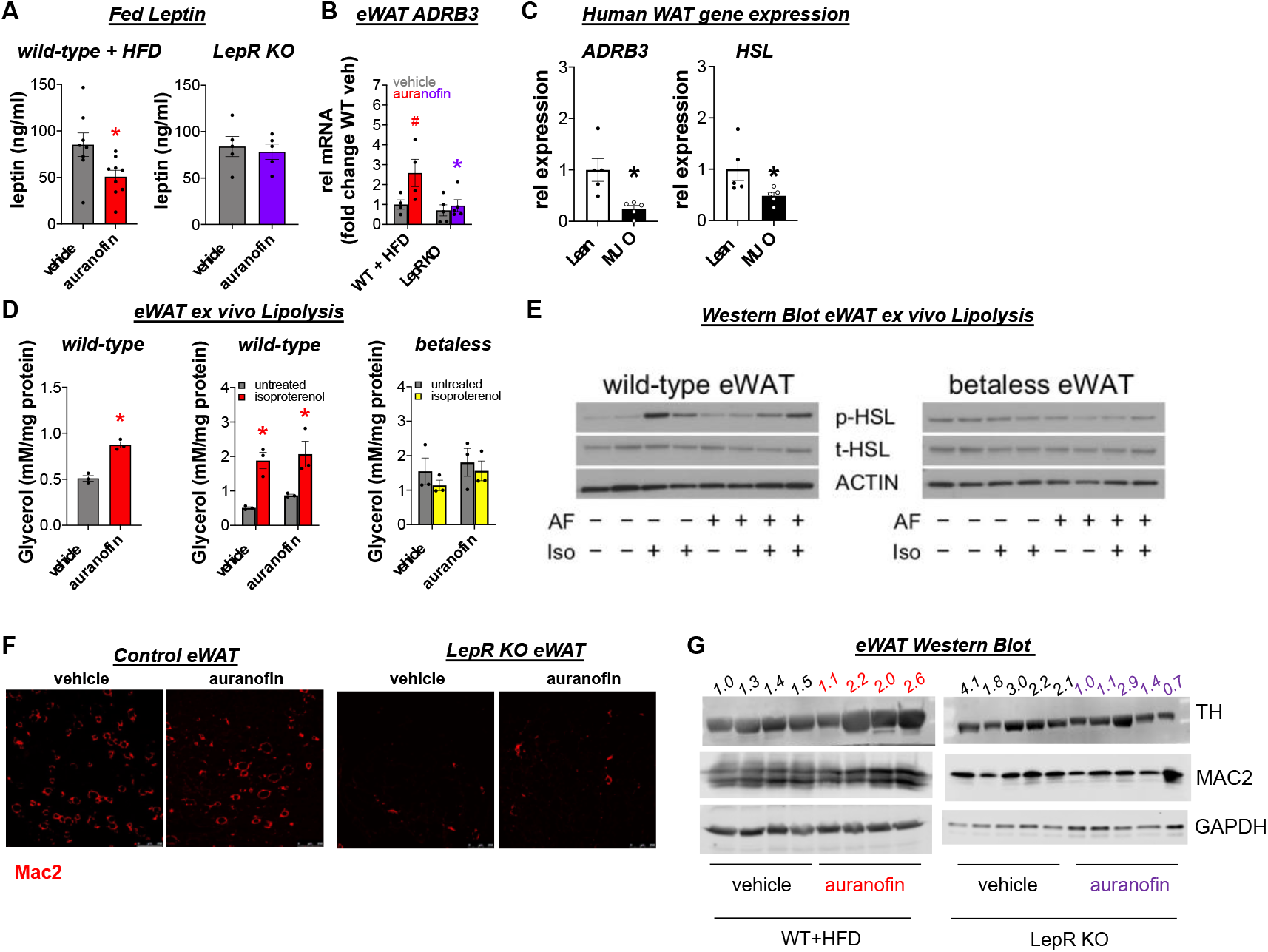
Auranofin enhances local leptin functions in eWAT independent of inflammation. Wild-type mice fed a high fat diet (HFD) for 12 weeks or *LepR KO* mice were i.p. injected with auranofin (1 mg/kg; red/purple) or vehicle (gray) for 4 weeks. **(A)** Fed serum leptin (ng/ml) wild-type (left; n=8—9/group) and *LepR KO* (right; n=5/group). *p=0.05 vs vehicle. **(B)** eWAT relative *Adrb3* expression (n=4-5/group). *p<0.05 vs wild-type, #p<0.10 vs vehicle. **(C)** Relative *ADRB3* and *HSL* expression measured in human subcutaneous adipose tissue biopsied from lean (white; n=5) and metabolically unhealthy obese (MUO, black; n=4) subjects. *p<0.05. **(D)** Glycerol release into the media two hours after stimulation (n=3/group). *p<0.05. eWAT tissues from wild-type and betaless (beta adrenergic receptor KO mice) were cultured *ex vivo* with auranofin (100 nM) or vehicle (DMSO) +/-isoproterenol (10 µM). **(E)** Immunoblots of eWAT lysates from **(D)** probed for the indicated proteins. **(F)** eWAT tissues stained for macrophages (Mac2; red); scale bar, 250 µm. **(G)** eWAT from **(F)** probed for the macrophage marker Mac2 and tyrosine hydroxylase (TH) to demonstrate auranofin heightens local leptin sensitivity. Data are represented as mean ± SEM.

The known influence of leptin on the sympathetic drive to eWAT raised the question of whether the effects of auranofin depend upon β-adrenergic signaling. Accordingly, we investigated the effects of auranofin on lipolysis in eWAT isolated from wild-type mice or beta-less mice, which lack all three β-adrenergic receptors (Bachman et al., 2002). Auranofin alone significantly increased glycerol release from wild-type eWAT compared to vehicle (DMSO) control **(Figure 6D)**. Treatment of wild-type eWAT with the β-adrenergic receptor agonist isoproterenol significantly stimulated glycerol release. Not surprisingly, beta-less eWAT was unresponsive to isoproterenol treatment, despite the presence of auranofin. Following the lipolysis studies, eWAT samples were subjected to immunoblotting to test activation of lipolysis (p-HSL) downstream of the beta-adrenergic receptor. As expected, p-HSL increased in wild-type eWAT exposed to isoproterenol, but no effects occurred in betaless eWAT **(Figure 6E)**. Interestingly, auranofin alone increased p-HSL levels consistent with increased glycerol release compared to vehicle control.

Macrophages reduce lipolytic responses in obese mice by interfering with sympathetic neurons in eWAT (Camell et al., 2017; Mowers et al., 2013). After establishing auranofin altered lipolysis, we asked whether macrophage content changed in the eWAT after auranofin using immunofluorescence labeling of Mac2. Although auranofin reduced markers of macrophage function associated with obesity, we found indistinguishable staining patterns between control and drug treatments **(Figure 6F)**. We also noticed *LepR KO* displayed considerably lower Mac2 labeling than wild-type mice suggesting any metabolic effects of auranofin on immune cells occur secondary to leptin lowering. Western blotting for tyrosine hydroxylase (TH) in the eWAT demonstrated the leptin reduction performed by auranofin in wild-type mice was required to increase sympathetic neuron content **(Figure 6G)** and contribute to greater lipolytic competence.

Overall, our experiments identified an insulin sensitizer that acts on the leptin-immune axis as a treatment for obesity and its comorbidities. Auranofin received FDA approval in 1985 to manage rheumatoid arthritis, an inflammatory condition often coupled with co-morbidities of T2DM (Giles et al., 2018). Our findings align with the metabolic impacts of other anti-inflammatory rheumatoid arthritis drugs, which improve insulin sensitivity in retrospective analyses (Collotta et al., 2020; Ursini et al., 2018).

In humans, auranofin reduces macrophage accumulation and pro-inflammatory cytokine expression, such as TNFa, in the lipophilic, synovial membrane of RA patients (Handel et al., 1995; Yanni et al., 1994). Owing to its hydrophobicity, auranofin accumulates in the lipophilic environment of WAT to diminish the expression of inflammatory genes and cytokines. As primary mechanisms of action, auranofin has been previously reported to exert multiple pleiotropic effects. Auranofin can induce reactive oxygen species (Nakaya et al., 2011) but also blunts pro-inflammatory signaling associated with poor WAT expandability (Saltiel and Olefsky, 2017). Although extremely high levels of ROS cause cellular damage, moderate ROS production may entrain a tolerable oxidative environment that enhances adipocyte differentiation without inflicting cellular damage (Han et al., 2016). The cell-specific effects of auranofin may also involve observed effects on NF-kB signaling (Asahara et al., 1995; Jeon et al., 2000) that culminates in reduced macrophage responses.

Inflammation also strongly correlates with hypoxia, fibrosis, and excessive collagen deposition, contributing to WAT metabolic dysfunction and fatty liver disease (Sun et al., 2013). WAT from insulin-sensitive obese individuals maintain oxygen partial pressure and express lower levels of hypoxia markers and fibrotic proteins compared to insulin-resistant obese individuals (Cifarelli et al., 2020). Likewise, auranofin suppressed Hypoxia-inducible factor 1-alpha protein levels and many fibrosis marker genes in eWAT that facilitate increased oxygenation, healthy WAT expansion, and insulin sensitivity (Halberg et al., 2009; Lee et al., 2014). Thus, the anti-inflammatory effects per se may occur secondary to other dominant pathogenic targets. Clinical studies support the idea the anti-fibrotic effects of auranofin have therapeutic relevance to metabolic health and may be uncoupled from broadly tempering macrophage accumulation in WAT (Beals et al., 2021; Cifarelli et al., 2020).

Despite durable correlations between inflammation and obesity in humans, causal relationships between obesity-mediated inflammation and insulin resistance remain unclear. Furthermore, recent provocative studies argue WAT inflammation responds to insulin resistance (Shimobayashi et al., 2018). Although our findings do not rule out the idea that anti-inflammatory monotherapies will be effective metabolic interventions, they argue strongly leptin lowering is a critical target to improve all the clinical phenotypes of obesity and T2DM.

Restoration of adrenergic (β-AR) receptor sensitivity in WAT enables lipolysis and body composition changes beneficial for diseases such as obesity. We propose that auranofin reduces leptin levels coupled with enhanced β-AR receptor regulation of lipolysis. While long considered an important component of thermogenesis (Collins et al., 2010; Himms-Hagen et al., 1994), neural inputs regulate β-AR signaling and lipolysis in WAT, serving a critical role to maintain insulin sensitivity and energy balance (Pirzgalska et al., 2017; Zeng et al., 2015). Interestingly, leptin elevates catecholamine (norepinephrine) content in WAT and activates lipogenesis through β-AR and p-HSL (Zeng et al., 2015). Auranofin increased β-AR sensitivity and p-HSL activation in eWAT that coincided with upregulated fatty acid metabolic gene program, which may sequester FFA from the liver in part to resolve steatosis. In obese humans, we show reduced expression of β-AR (*ADRB3*) and *HSL/LIPE*, which likely impairs catecholamine signaling and lipolysis. Leptin levels remained high in *LepR KO* mice, and auranofin does not elevate expression of lipolytic genes nor increase TH abundance. Our observations add to recent studies (Zhao et al., 2020; Zhao et al., 2019) and suggest the metabolic benefits auranofin of leptin lowering also restore lipolytic competence in WAT.

One strength of our study is we performed experiments at auranofin doses lower than the human bioequivalent and showed auranofin not only improves glycemia, but the overall metabolic phenotype of obesity, including insulin sensitivity, normalization of serum insulin and leptin levels, and reduction of ectopic lipid accumulation in the liver. Indirect calorimetry studies showed no change in metabolic fuel selection. Our treatment regimen does not reduce body weight or lower food intake but promotes locomotor activity derived from peripheral leptin functions. Leptin also stimulates locomotor activity in obese mice and skeletal muscle glucose uptake, independent of food intake and body weight (Huo et al., 2009). Similarly, auranofin increased locomotor activity, likely involving increased skeletal muscle glucose uptake independent of canonical insulin signals like Akt (Saltiel, 2021). Along these lines, it will now be essential to leverage tracer methods and clinical studies to uncover how auranofin and leptin lowering impact activity, muscle metabolism, and glucose disposal at a physiologic and molecular level.

Auranofin has been clinically used to treat rheumatoid arthritis patients for years, so the human dosing and toxicological data in mice are known and abundantly available. To date, fourteen other clinical trials tested the clinical efficacy of auranofin in cancer, HIV, and other infectious diseases (https://www.clinicaltrials.gov). Yet, none of the studies monitored glucose or other energy balance variables as outcomes. The scarcity of information related to metabolic hormones when individuals are on rheumatoid arthritis drugs stems from the fact that the patients also receive other therapies in combination known to interfere with blood sugar, including glucocorticoids and methotrexate. It is unclear how efficacious auranofin might be for individuals with obesity and T2DM in the clinic. Therefore, a controlled clinical study for auranofin with a primary endpoint of insulin sensitivity (e.g., hyperinsulinemic-euglycemic clamp) is warranted. This new discovery is of crucial importance given current proposals to repurpose drugs for treating obesity and its co-morbidities.

## LIMITATIONS OF THE STUDY

Generally, the therapeutic mechanisms of auranofin and gold salts remain unclear, and no unifying theory has been proposed so far to our knowledge despite the high number of targets and mechanisms described. Although we ruled out *miR-30a* as the rate-limiting factor for auranofin effects, the mechanism of action for auranofin’s metabolic effects is not clear. We did not assess whether auranofin affects nutrient absorption, which could explain the general maintenance of body weight despite higher home cage activity. It is, of course, possible that auranofin acts on the skeletal muscle to improve insulin action but the accumulation in the musculoskeletal system seems nominal and unlikely (Gottlieb et al., 1972). Another limitation of our study is that the drug has not been tested in female mouse models of obesity and a complete lack of leptin causes infertility. Based on a recent study (Zhao et al., 2019), we expect leptin lowering by auranofin or other similar approaches to spare fertility. Expanding the effects of auranofin to female mice is a future goal but challenging because sex influences sensitivity to HFD (Corrigan et al., 2020). Lastly, we are excited by the anti-inflammatory effects of auranofin but believe that the anti-diabetic actions of leptin lowering are probably more relevant than the impact on WAT inflammation. In this regard, our study fills a gap in the literature by demonstrating how local leptin actions in eWAT improve insulin sensitivity without diminishing inflammatory tone.

## ACKNOWLEDGEMENTS

This work was funded by American Diabetes Association #1-18-IBS-105 (S.M.H.) and NIH grants R01DK114356 (S.M.H.), R01DK126042 (S.M.H.), and R01DK121348 (H.W.). This study was also funded (in part) by an award from the Baylor College of Medicine Nutrition and Obesity Pilot and Feasibility Fund and the BCM Bridge to Independence Program (A.R.C). The Reverse Phase Protein Array Core is supported by the CPRIT Core Facility Support Award RP170005 “Proteomic and Metabolomic Core Facility,” NCI Cancer Center Support Grant P30CA125123, and intramural funds from the Dan L. Duncan Cancer Center. Other core services at BCM utilized in this project were supported with funding from NCI P30CA125123: Genetically Engineered Rodent Models Core, Human Tissue Acquisition and Histology Core, and the Integrated Microscopy Core. This study was also supported, in part, by the Assistant Secretary of Defense for Health Affairs endorsed by the DOD PRMRP Discovery Award (No. W81XWH-18-1-0126 to K.H.K.).

## AUTHOR CONTRIBUTIONS

A.R.C. and S.M.H. conceptualized the study. A.R.C, S.M.H., and the remaining authors performed critical experiments in mice and/or contributed to the interpretation of metabolic phenotypes and biochemical outcomes. A.R.C. and S.M.H. wrote the manuscript with editorial input from all authors. S.M.H. is the guarantor of this work and, as such, had full access to all the data in the study and takes responsibility for the integrity of the data and the accuracy of the data analysis.

## STAR METHODS

### RESOURCE AVAILABILITY

#### Lead Contact

Further information and requests for resources should be directed to and will be fulfilled by the Lead Contact, Sean M. Hartig. All data that support the findings herein presented are available from the corresponding author upon reasonable request.

#### Materials Availability

Resources (*miR-30a*^*-/-*^ mice) generated during the current study are available from the Lead Contact upon reasonable request.

#### Data and Code Availability

All data generated or analyzed during this study are included in the published article. RNA-seq datasets are deposited in Gene Expression Omnibus.

### EXPERIMENTAL MODEL AND SUBJECT DETAILS

#### Mice and housing conditions

All animal procedures were approved by the Institutional Animal Care and Use Committee of Baylor College of Medicine. All experiments were conducted using littermate-controlled male mice aged 6-8 weeks. All mice were housed in a barrier-specific pathogen-free animal facility with 12h dark-light cycle and free access to water and food. C57BL/6J wild-type mice and FVB mice (control mice for beta-less experiments) were obtained from the BCM Center for Comparative Medicine. Mice were fed 60% high-fat diet (HFD; Bio-Serv) for 12 weeks before experiments. *Ubc-Cre;LepR*^*lp/lp*^ (*LepR KO*) mice were generated previously (Cox et al., 2016). Male *LepR KO* mice were treated with 0.1 mg/g body weight by oral gavage (Cox et al., 2016) at 8 weeks of age to induce *LepR* gene deletion. Beta-less mice (*Adbr1, Adbr2, Adrb3* triple knockout) were bred in-house to generate 3-month-old mice for *in vitro* studies. Mice were injected with auranofin (1, 5, 7.5, or 10 mg/kg body weight) or vehicle control (5% DMSO, 5% polyethylene glycol, 10% ethanol, PBS) three days per week for 4 weeks, followed by metabolic phenotyping. At the end of experiments, mice were euthanized by cervical dislocation while under isoflurane anesthesia. After euthanasia, tissues were collected, flash-frozen in liquid N_2_, and stored at -80°C until use. All experiments adhered to ARRIVE Guidelines.

#### Human adipose tissue

Subcutaneous WAT RNA samples were obtained from Dr. Samuel Klein (Washington University). Sample collection was described previously (Cifarelli et al., 2020) and used the following inclusion criteria for each of the 2 groups: (a) metabolically healthy lean had BMI 18.5–24.9 kg/m^2^, plasma TG concentration <150 mg/dL, fasting plasma glucose concentration <100 mg/ dL, 2-hour OGTT plasma glucose concentration <140 mg/dL, HbA1c ≤5.6%, and IHTG content <4%; (b) metabolically unhealthy obese (MUO) had BMI 30–49.9 kg/m^2^, prediabetes (fasting plasma glucose concentration ≥100 mg/dL, 2-hour OGTT plasma glucose concentration ≥140 mg/dL, and/or HbA1c ≥5.7%), and IHTG content ≥5%.

#### Generation of *miR-30a*^*-/-*^ mice

*miR-30a*^*-/-*^ mice were generated with the BCM Genetically Engineered Rodent Models Core. Single guide RNAs (sgRNA) flanked a 1 kb region on mouse chromosome 1 (mm10_dna range=chr1:23272064-23272543) that contains *miR-30a*. sgRNA sequences were designed to target the genomic sequences using upstream sgRNA, 5′-ACTCCCGCAGAGCACTTCTCAGG-3′ and downstream sgRNA, 5′-TCAAGGAGTAATTATCTTGTTGG-3′. To minimize the probability of off-target events, only sgRNAs predicted to have off-target sites with three mismatches or more were used to target Cas9 endonuclease activity to regions flanking the *miR-30a* gene. Two hundred C57BL/6NJ pronuclear-stage zygotes were co-injected with 100ng/μl Cas9 mRNA, 20ng/μl sgRNA (each), and 100ng/μl of ssODNs. Following microinjection, zygotes were transferred into pseudopregnant ICR recipient females at approximately 25–32 zygotes per recipient. Sanger sequencing and founder line genotyping from mouse genomic DNA confirmed the deletion. DNA extracted from mouse ear clips was used in PCR reactions with primers designed to detect the deleted alleles. Primer sequences (5’ ->3’) that flanked the *miR-30a* locus for genotyping were: GCATCGAGGCTTTGCAGTTT and TGCACAGGAAGAACACTTCTGT.

The TaqMan Advanced miRNA cDNA Synthesis Kit (ThermoFisher, #A28007) was used to synthesize miRNA cDNA from 20 ng total RNA. To extend mature miRNAs, polyadenylation and adaptor sequence ligation of the 3’ and 5’ ends, respectively, occur before universal priming and reverse transcription. To address low-expressing targets, cDNA is amplified by primers that recognize sequences appended to both ends, effectively minimizing amplification bias. Next, the TaqMan Advanced miRNA Assays (ThermoFisher #A25576) were used to quantify relative gene expression. As recommended by the manufacturer, invariant RNA controls included *miR-423-3p, miR-451*, and *miR-423-5p*.

#### Glucose and insulin tolerance tests

To determine glucose tolerance, mice were fasted 16 hours, and glucose was administered (1.5 g/kg (HFD) or 0.75 mg/kg (normal chow) body weight) by intraperitoneal injection (IP). To determine insulin tolerance, mice were fasted four hours before insulin IP (1.5 U/kg body weight). Blood glucose levels were measured by handheld glucometer. ELISA quantified serum hormone levels for overnight fasting insulin (Millipore #EZRMI-13K) and fed leptin (Crystal Chem #90030) and soluble DPP4 (Thermo Fisher #EMDPP4). Mouse body composition was examined by MRI (Echo Medical Systems).

#### Indirect calorimetry

Mice were maintained on HFD and housed at room temperature in Comprehensive Lab Animal Monitoring System Home Cages (CLAMS-HC, Columbus Instruments). Oxygen consumption, CO_2_ emission, energy expenditure, food and water intake, and activity were measured for seven days (BCM Mouse Metabolic Phenotyping Core). Mouse body weight was recorded, and body composition was examined by MRI (Echo Medical Systems) prior to indirect calorimetry.

#### Immunoblotting

Tissue lysates were prepared in Protein Extraction Reagent (Thermo Fisher #78510) supplemented with Halt Protease and Phosphatase Inhibitor Cocktail (Thermo Fisher #78440). Immunoblotting was performed with lysates run on 4-12% Bis-Tris NuPage gels (Life Technologies #NP0321Box) and transferred onto Immobilon-P Transfer Membranes (Millipore #IPVH00010) followed by antibody incubation. Immunoreactive bands were visualized by chemiluminescence.

#### qPCR

Total RNA was extracted using the Direct-zol RNA MiniPrep kit (Zymo Research #R2051). cDNA was synthesized using qScript (QuantBio #95048-100). Relative mRNA expression was measured with SsoAdvanced Universal Probes Supermix reactions (Bio-Rad #175284) read out with a QuantStudio 3 real-time PCR system (Applied Biosystems). TATA-box binding protein (*Tbp)* was the invariant control. Roche Universal Probe Gene Expression Assays were used as previously described (Koh et al., 2018). Table S2 lists the primers and qPCR assays used in this study.

#### RNA-seq

Snap frozen tissues were sent to Novogene for RNA extraction and sequencing. Sample quality control was analyzed on Bioanalyzer 2100 (Agilent) followed by mRNA library prep using NEBNext Ultra II kit (non-stranded, poly-A selected). The prepared libraries were quality controlled and sequenced using PE150 on the Illumina NovaSeq 6000, generating ~20 million paired reads/sample. Reads were mapped to the mouse reference genome mm10 using STAR software. Differential expression analysis of auranofin versus vehicle treatment was performed using the DESeq2 R package. The resulting p values were adjusted using Benjamini and Hochberg’s approach for controlling the false discovery rate. Gene set enrichment analysis was performed with the Molecular Signatures Database, and normalized enrichment scores were calculated for Hallmark gene sets. RNA-seq files can be accessed through Gene Expression Omnibus.

#### Microscopy and histology

Mitochondria were labeled using MitoTracker CMX-ROS (ThermoFisher). Live cells were pulsed with 500 nM MitoTracker for 15 min. Mitochondrial labeling was followed by cell fixation in 4% paraformaldehyde. Ammonium chloride was used to quench auto-fluorescence derived from residual paraformaldehyde. DAPI (Sigma) and LipidTOX were used for nuclei and lipid droplet labeling, respectively. Imaging was performed with the DeltaVision Core Image Restoration Microscope (GE Healthcare). Formalin-fixed paraffin-embedded adipose and liver tissue sections were stained with hematoxylin and eosin (H/E) and Oil Red O stains, respectively, by the BCM Human Tissue Acquisition and Pathology Core. Four 20x fields of view per tissue were imaged using a Nikon Ci-L Brightfield microscope. Fiji software quantified adipocyte morphometry in histological sections of WAT (Galarraga et al., 2012).

#### Flow cytometry

Stromal vascular fraction (SVF) cells were isolated from eWAT following digestion with collagenase type I (Worthington Biochemical Corporation). Cells were stained with Fc receptor blockade and antibodies to immune cell markers CD45 (Thermo Fisher #17-0451-82), F4/80 (Thermo Fisher #45-4801-82), CD11c (Thermo Fisher #12-0114-82), and CD206 (Bio-Rad Laboratories #MCA2235F) or respective isotype controls. Data were collected with an LSRII cytometer (BD Biosciences) and analyzed using Kaluza software (Beckman Coulter, Indianapolis, IN). Gating was performed on viable CD45+ cells, from which total macrophages were identified as F4/80+ high. Macrophages were classified into CD11c+CD206–/low “M1-like” macrophages and CD206+/CD11c– “M2-like” macrophages. Data were presented as cell numbers per gram adipose tissue calculated using counting beads (Biolegend, San Diego, CA) or as percentages of cell subsets of total macrophages.

#### Mass spectrometry

We analyzed auranofin biodistribution with the NMR and Drug Metabolism Core at Baylor College of Medicine using previously established protocols (York et al., 2017). Briefly, plasma and tissue extracts were homogenized in ice-cold MeOH solutions. After vortexing and centrifugation (15 min at 15,000 × g), each supernatant was transferred to an autosampler vial, and 5.0 μl was injected into a system combining ultra-high performance liquid chromatography (UHPLC) and quadrupole time of flight mass spectrometry (QTOFMS) for analysis. The concentration of auranofin was calculated based on standard curves in the corresponding matrices. The linear trapezoidal rule was used for area under the curve calculations.

#### Proteomics

Protein lysates were prepared by the BCM Antibody-Based Proteomics Core for reverse phase protein array assays. The Aushon 2470 Arrayer (Aushon BioSystems) with a 40 pin (185 µm) configuration was used to spot lysates onto nitrocellulose-coated slides (Grace Bio-Labs). The slides were probed with 220 antibodies against total and phosphoprotein proteins using an automated slide stainer (Dako). Primary antibody binding was detected using a biotinylated secondary antibody followed by streptavidin-conjugated IRDye 680 fluorophore (LI-COR Biosciences). Fluorescent-labeled slides were scanned on a GenePix AL4200, and the images were analyzed with GenePix Pro 7.0 (Molecular Devices). The images were analyzed with GenePix Pro 7.0 Microarray Acquisition & Analysis Software (Molecular Devices). Background subtracted total fluorescence intensities of each spot were normalized for variation in total protein (Sypro Ruby) and nonspecific labeling. For adipokine analysis, tissue lysates from four samples were pooled (125 µg/sample) and incubated with membranes from the Proteome Profiler Mouse Adipokine Array Kit (R&D Systems #ARY013) according to the manufacturer’s instructions.

#### In vitro experiments

SVF cells were isolated from mouse inguinal WAT. Fat depots were digested in PBS containing collagenase I (Roche, 1.5 U/ml) and dispase II (Sigma, 2.4 U/ml) supplemented with ten mM CaCl_2_ at 37°C for 40-45 min. The primary cells were filtered twice through 70 µm cell strainers and centrifuged at 700 rcf to collect the SVF. The SVF cell pellets were rinsed and plated. Adipocyte differentiation was induced by treating confluent cells in DMEM/F12 medium containing Glutamax (ThermoFisher), 10% FBS, 0.250 mM isobutylmethylxanthine (Sigma Chemical Co.), 1 mM rosiglitazone (Cayman Chemical Co.), 1 mM dexamethasone (Tocris Biosciences), 850 nM insulin (Sigma), and 1 nM T3 (Sigma). Four days after induction, cells were switched to the maintenance medium containing 10% FBS, 1 mM rosiglitazone, 1 mM dexamethasone, 850 nM insulin, 1 nM T3. Experiments occurred 8-10 days after induction of differentiation.

#### Ex vivo lipolysis

Fat pads were excised and cut into ~50 mg aliquots in PBS on ice. At least two tissue aliquots per mouse were allocated to each treatment group. Each tissue was placed in a well of a 24-well plate containing 0.5 ml Krebs buffer and incubated for 1 h at 37°C, 5% CO_2_. Media was replaced with fresh Krebs buffer containing 100 nM auranofin or vehicle (DMSO) +/-10 µM isoproterenol. After a 2 h incubation at 37°C, 5% CO_2_, tissues were snap-frozen in liquid N_2_ and media snap-frozen on dry ice. Lysates were made from tissues for protein quantification for normalization of glycerol release. Glycerol was measured by colorimetric assay (Sigma #MAK117).

#### Seahorse assays

Respiration was measured in adipocytes using an XF24 analyzer (Agilent). SVF cells were plated into V7-PS plates and grown to confluence. Cells were differentiated for 8-10 days before treatment with auranofin (1000 nM) or vehicle (DMSO) overnight. For the assay, media was replaced with 37 °C unbuffered DMEM containing 4.5 g/L glucose, sodium pyruvate (1 mmol/L), and L-glutamine (2 mmol/L). Measurements were made at 37 °C using 2-2-2 intervals. Basal respiration was defined before sequential addition of oligomycin, FCCP, rotenone, and antimycin A.

### QUANTIFICATION AND STATISTICAL ANALYSIS

All data are presented as mean ± SEM, unless otherwise specified. All statistical analyses were conducted in Prism 9 (GraphPad). We used unpaired two-tailed t-tests, Mann-Whitney U tests, and one-way and two-ANOVAs to compare means of numerical variables where appropriate. Embedded ANCOVA tools in CalR were used to determine significant changes in energy expenditure measurements collected from CLAMS cages. The statistical details of experiments can be found in the figure legends. No statistical method was used to predetermine sample size. Our primary threshold for statistical significance was p<0.05. Unblinded analysis of histology and immunohistochemistry was performed by the investigators.

---

**Supplemental Figure 1.**
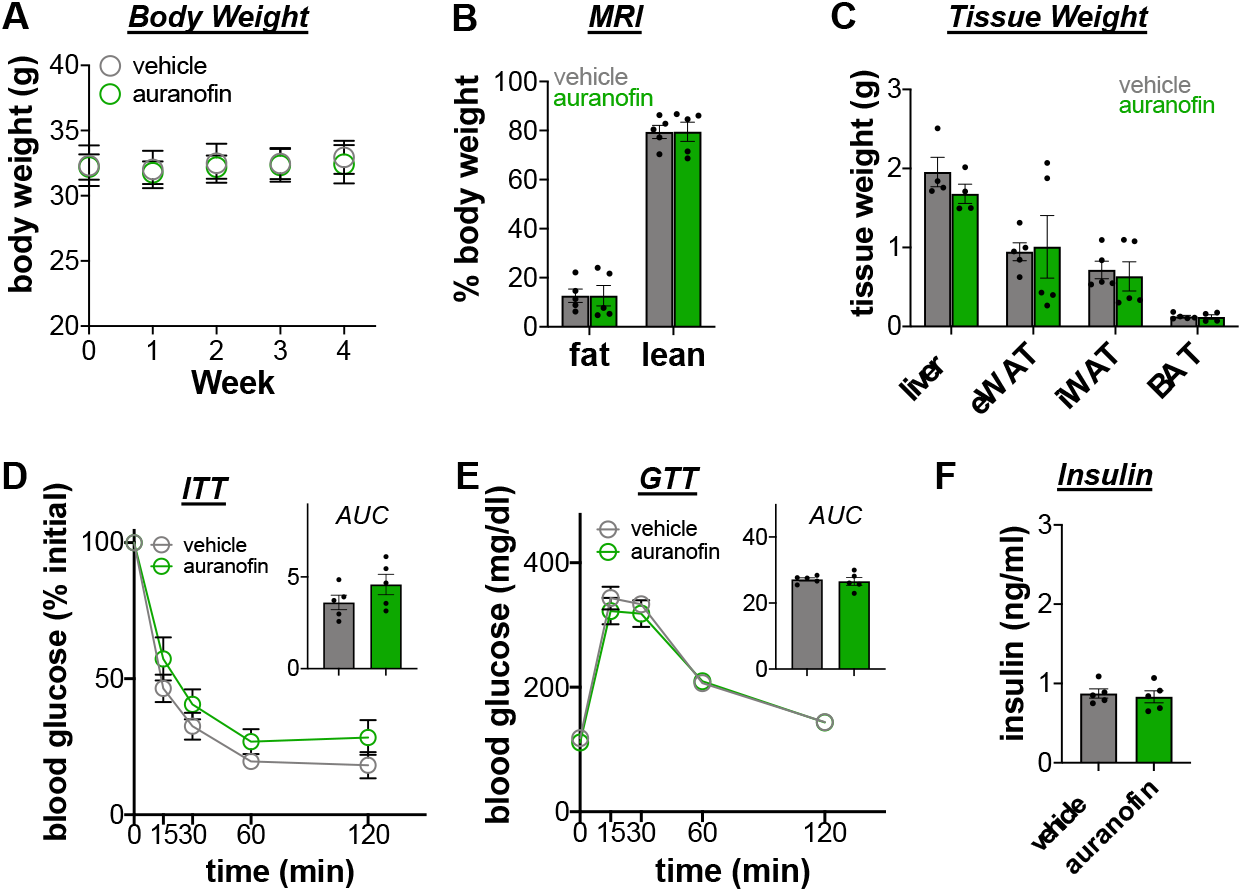
The metabolic effects of auranofin require obesity. Mice fed a normal chow diet (NC) were i.p. injected with auranofin (1 mg/kg; green) or vehicle (gray) for 4 weeks starting at 18 weeks of age (n=5/group). **(A)** Body weight during treatment with **(B)** final body composition (% body mass). **(C)** Tissue weights (g). **(D)** Insulin (ITT) and **(E)** glucose (GTT) tolerance tests, with corresponding area under the curve (AUC) measurements (×10^4^ or ×10^5^, respectively). **(F)** Overnight fasting serum insulin (ng/ml). Data are represented as mean ± SEM.

**Supplemental Figure 2.**
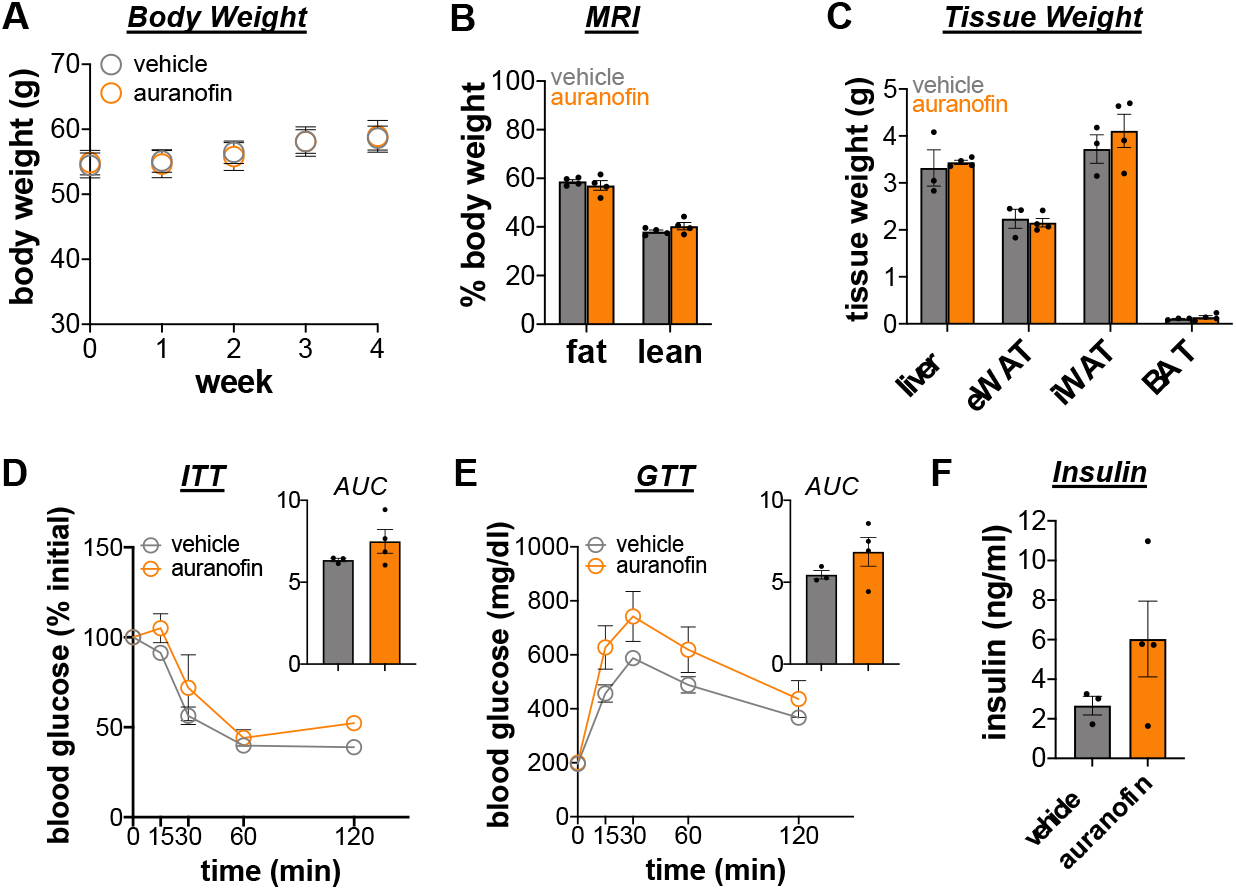
Leptin deficient mice are resistant to the insulin sensitizing effects of auranofin. Male ob/ob mice were i.p. injected with auranofin (orange) or vehicle (gray) for 4 weeks starting at 18 weeks of age (n=3-4/group). **(A)** Body weight during treatment with **(B)** final body composition (% body mass). **(C)** Tissue weights (g). **(D)** Insulin (ITT) and **(E)** glucose (GTT) tolerance tests, with corresponding area under the curve (AUC) measurements (×10^4^ or ×10^5^, respectively). **(F)** Overnight fasting serum insulin (ng/ml). Data are represented as mean ± SEM.

